# Index switching causes “spreading-of-signal” among multiplexed samples in Illumina HiSeq 4000 DNA sequencing

**DOI:** 10.1101/125724

**Authors:** Rahul Sinha, Geoff Stanley, Gunsagar S. Gulati, Camille Ezran, Kyle J. Travaglini, Eric Wei, Charles K.F. Chan, Ahmad N. Nabhan, Tianying Su, Rachel M. Morganti, Stephanie D. Conley, Hassan Chaib, Kristy Red-Horse, Michael T. Longaker, Michael P. Snyder, Mark A. Krasnow, Irving L. Weissman

**Affiliations:** Institute for Stem Cell Biology and Regenerative Medicine, Stanford University School of Medicine, 265 Campus Drive, Stanford, CA 94305, USA; Department of Bioengineering, Stanford University School of Engineering, 443 Via Ortega, Stanford, CA 94305, USA; Department of Biochemistry and Howard Hughes Medical Institute, Beckman Center B400, 279 Campus Drive, Stanford, CA 94305-5307, USA; Stanford Center for Genomics and Personalized Medicine, 3165 Porter Drive, Palo Alto, CA 94304, USA; Department of Surgery, Hagey Laboratory for Pediatric Regenerative Medicine, 257 Campus Drive, Stanford, CA 94305, USA; Department of Biology, Gilbert Building, Rm 109, 371 Serra Mall, Stanford, CA 94305, USA; Department of Pathology, Stanford University School of Medicine, 300 Pasteur Drive, Stanford, CA 94305, USA; Ludwig Center for Cancer Stem Cell Research and Medicine, Stanford University School of Medicine, 265 Campus Drive, Stanford, CA 94305, USA

**Author notes:** These authors contributed equally to this work. Correspondence and requests for materials should be addressed to R.S.

## Abstract

Illumina-based next generation sequencing (NGS) has accelerated biomedical discovery through its ability to generate thousands of gigabases of sequencing output per run at a fraction of the time and cost of conventional technologies. The process typically involves four basic steps: library preparation, cluster generation, sequencing, and data analysis. In 2015, a new chemistry of cluster generation was introduced in the newer Illumina machines (HiSeq 3000/4000/X Ten) called exclusion amplification (ExAmp), which was a fundamental shift from the earlier method of random cluster generation by bridge amplification on a non-patterned flow cell. The ExAmp chemistry, in conjunction with patterned flow cells containing nanowells at fixed locations, increases cluster density on the flow cell, thereby reducing the cost per run. It also increases sequence read quality, especially for longer read lengths (up to 150 base pairs). This advance has been widely adopted for genome sequencing because greater sequencing depth can be achieved for lower cost without compromising the quality of longer reads. We show that this promising chemistry is problematic, however, when multiplexing samples. We discovered that up to 5-10% of sequencing reads (or signals) are incorrectly assigned from a given sample to other samples in a multiplexed pool. We provide evidence that this “spreading-of-signals” arises from low levels of free index primers present in the pool. These index primers can prime pooled library fragments at random via complementary 3’ ends, and get extended by DNA polymerase, creating a new library molecule with a new index before binding to the patterned flow cell to generate a cluster for sequencing. This causes the resulting read from that cluster to be assigned to a different sample, causing the spread of signals within multiplexed samples. We show that low levels of free index primers persist after the most common library purification procedure recommended by Illumina, and that the amount of signal spreading among samples is proportional to the level of free index primer present in the library pool. This artifact causes homogenization and misclassification of cells in single cell RNA-seq experiments. Therefore, all data generated in this way must now be carefully re-examined to ensure that “spreading-of-signals” has not compromised data analysis and conclusions. Re-sequencing samples using an older technology that uses conventional bridge amplification for cluster generation, or improved library cleanup strategies to remove free index primers, can minimize or eliminate this signal spreading artifact.

## Introduction

Since the first draft of the human genome was published in 2003, high-throughput sequencing (HTS) methods have largely replaced the conventional (Sanger) sequencing methods as integral tools for biological research. Of all the different HTS approaches, Illumina’s “sequencing by synthesis” platform is the most widely adopted due to its unmatched throughput, scalability, and speed—all of which result in significant cost reduction. Further cost reduction is achieved by multiplexing—a procedure where multiple DNA samples, each containing millions of distinct DNA molecules can be mixed together, sequenced in a massively parallel fashion, and each resulting read then reassigned to its respective sample. Reassignment is performed computationally based on unique molecular indices that are shared by all DNA fragments within a sample. These indices are short stretches of unique DNA sequence (“barcodes”) that are introduced by DNA ligation or PCR during sample preparation.

The reduction in cost by multiplexing allows greater flexibility in the design of experiments investigating 1) mutations or single nucleotide polymorphisms (SNPs) by whole genome sequencing (WGS) or more targeted exome-sequencing, 2) transcriptome profiling (RNA-seq), 3) DNA-protein interactions (ChIP-seq), and 4) epigenome characterization by ATAC-seq or bisulfite sequencing. These methods rely on the faithful reassignment of each DNA sequencing read to the appropriate sample after sequencing.

Here we describe a problem with the newer iteration of Illumina sequencers, HiSeq 4000, which results in inaccurate reassignment of reads by switching or replacement of the unique molecular indices present at the ends of each DNA library fragment. This occurs before sequencing when free index primers, each carrying a unique index, are present in the mixed (multiplexed) library pool. Such free index primers persist at low levels after the most commonly used library purification protocol recommended by Illumina, which uses AMPure XP beads—the standard method in the sequencing field for almost eight years^1,2^. This “spreading-of-signals” issue is not found in the previous iteration of Illumina sequencer, NextSeq 500, even when high levels of index primers are present in a mixed library pool.

We hypothesize that this replacement, or index-switching, on the HiSeq 4000 occurs during the exclusion amplification (ExAmp) procedure^3^ necessary for cluster generation on the flow cell before sequencing by the HiSeq 4000. The ExAmp step is not necessary for cluster generation on NextSeq 500, which uses conventional bridge amplification^4^ to generate clusters on a NextSeq 500 flow cell.

Bridge amplification is the original Illumina (Solexa) clustering mechanism^5^. It relies on single-stranded library molecules binding to an oligonucleotide (P5 primer) immobilized on a glass flow cell surface. Library molecules that do not bind to an immobilized oligonucleotide are washed from the flow cell along with any leftover primers. All DNA amplification steps take place after the wash step. The fixed oligonucleotide acts as a primer for a DNA polymerase to copy the hybridized strand, resulting in a covalently attached complementary sequence of the DNA library molecule. The covalently linked complementary single-stranded library molecule contains sequences at the distal end that are complementary to a second oligonucleotide (P7 primer), which is also fixed to the flow cell surface. The distal end of the covalently linked library molecule hybridizes to the P7 primer, forming an arching bridge. Using P7 as a primer, a new copy of the original library molecule is synthesized using the fixed complementary strand as a template, and becomes covalently linked to the flow cell by this process. This allows for bridge PCR amplification of the library molecule with both ends attached to the flow cell surface, using multiple cycles of denaturation and replication to form a cluster for sequencing.

The ExAmp procedure for cluster generation in the HiSeq 4000 is a fundamental shift from the older method. The details of this new chemistry are proprietary, and the only insight comes from patent descriptions^3,6^. ExAmp does not include the regular bind-and-wash steps prior to cluster generation as described above. Instead, the single stranded library molecules—resulting from the alkali denaturation of double-stranded library molecules—are mixed with the ExAmp clustering reagents (Illumina) and loaded onto a patterned flow cell^7^. The ExAmp chemistry involves a rapid isothermal amplification step necessary for cluster generation **(Supplementary Fig. 1)**.

We show that the index-switching described above during the ExAmp procedure leads to spreading of signal among samples that were multiplexed. Most likely this index-switching happens during seeding and before cluster generation, because all the active reagents (in particular DNA polymerase) necessary for cluster generation are present in the reaction mix while the library molecules are being guided to the nanowells to seed a cluster. If free index primers are present during this procedure they can prime the library fragments and get extended by the active DNA polymerase, forming a new library molecule with a different index. These new library molecules with switched identities can seed free nanowells and generate clusters, thus resulting in reads that get falsely assigned to a different sample **(Fig. 1)**.

**Figure 1.**
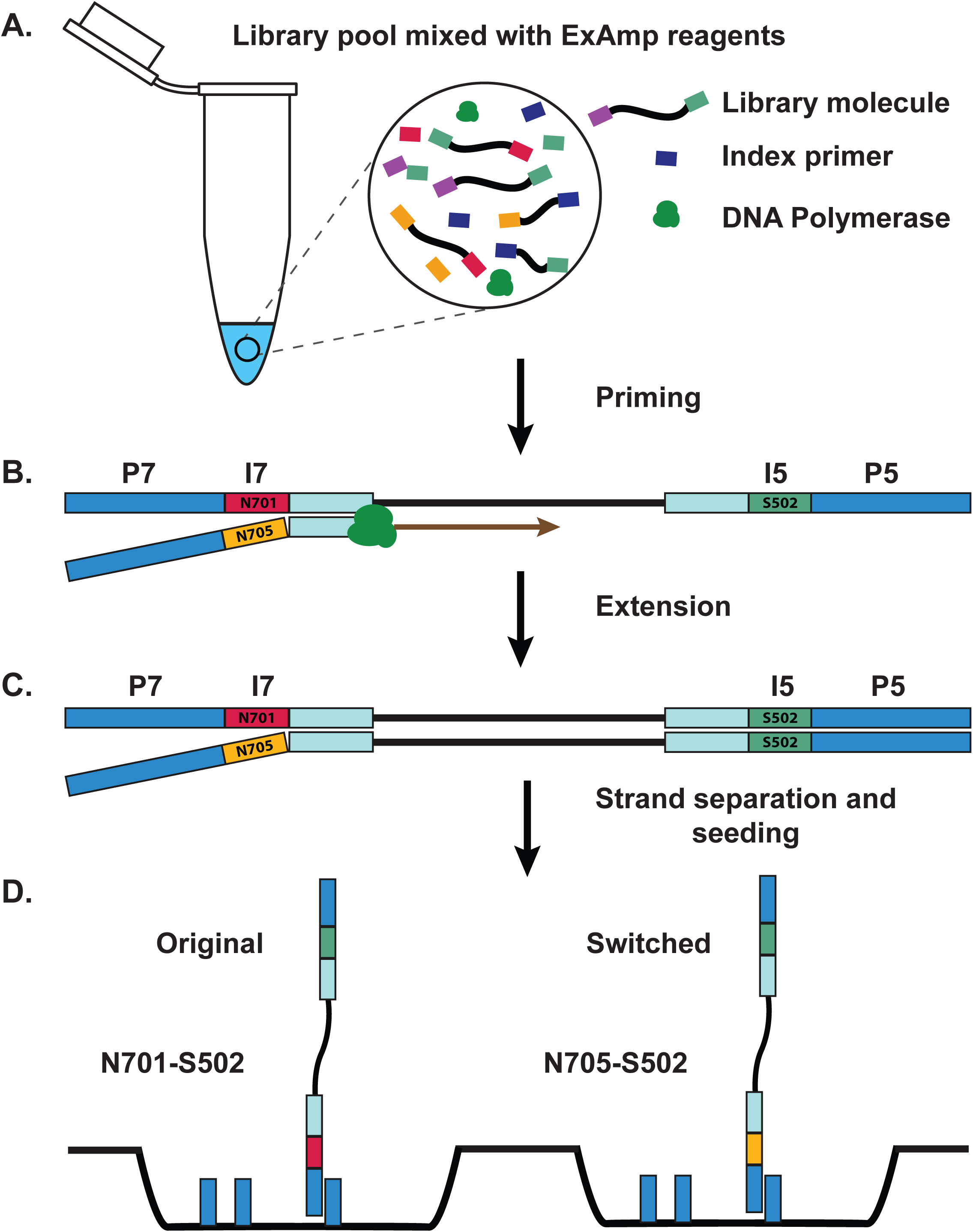
Mechanism for ExAmp index switching. A) Library pool with residual free index primers is mixed with ExAmp clustering reagents before loading on a patterned flow cell. B) A free index primer anneals to an existing library molecule and C) gets extended by active DNA polymerase to form a new library molecule with switched index. D) The newly synthesized strand separates from the template and seeds a nanowell on a patterned flow cell to generate a cluster.

## Methods

We used Smart-seq2^8^ to perform single-cell RNA sequencing, with the goal of using RNA expression patterns to more precisely identify and characterize subpopulations of mouse hematopoietic stem cells (mHSCs) and mouse fetal heart cells (mFHCs). In brief, using the BD FACS Aria II system (BD Biosciences), we sorted either highly purified mHSCs or mFHCs into 96-well full-skirted PCR plates (BioRad), where each well contained 4 μl of lysis buffer **(Fig. 2)**. Poly-A mRNA in the cell lysate was converted to cDNA and amplified as described^8^ **(Fig. 2A)**. Amplified cDNA in each well was quantified using a high-throughput Fragment Analyzer (Advanced Analytical) **(Fig. 2B).** After quantification, cDNA from each well was normalized to the desired concentration range (0.05 ng/μL − 0.16 ng/μL) by dilution, consolidated into a 384-well plate, and subsequently used for library preparation (Nextera XT kit; Illumina) using a semiautomated pipeline as described^9,10^ **(Fig. 2C).** The distinct libraries resulting from each well were pooled, cleaned-up and size-selected using precisely 0.6x volumes (mHSCs) or 0.7x volumes (mFHCs) of Agencourt AMPure XP beads (Beckman Coulter), as recommended by the Nextera XT protocol (Illumina). Use of 0.6x-0.8x volume of precalibrated AMPure XP beads is a standard cleanup procedure for various kinds of libraries to exclude DNA fragments smaller than 500 base pairs (bp) (usually 0.6x volume) or 300 bp (usually 0.8x volume) **(Supplementary Fig. 2)**. A high-sensitivity Bioanalyzer (Agilent) run was used to assess fragment distribution and concentrations of different fragments within the library pool **(Fig. 4B)**. Library pools were then loaded on either a HiSeq 4000 or a NextSeq 500 at two different core facilities at Stanford University. It is important to note that after pooling the libraries and before sequencing there is no PCR step in our protocol. The resulting reads were 1) demultiplexed using Illumina’s demultiplexing tool bcl2fastq (default settings), and 2) processed using skewer^11^ for 3’ quality-trimming, 3’ adapter-trimming, and removal of degenerate reads, as described^9^. The processed reads were mapped to the mouse genome (mm10) using OLego^12^, and the mapped reads were plotted in the 384-well format depicting number of mapped reads per well.

**Figure 2.**
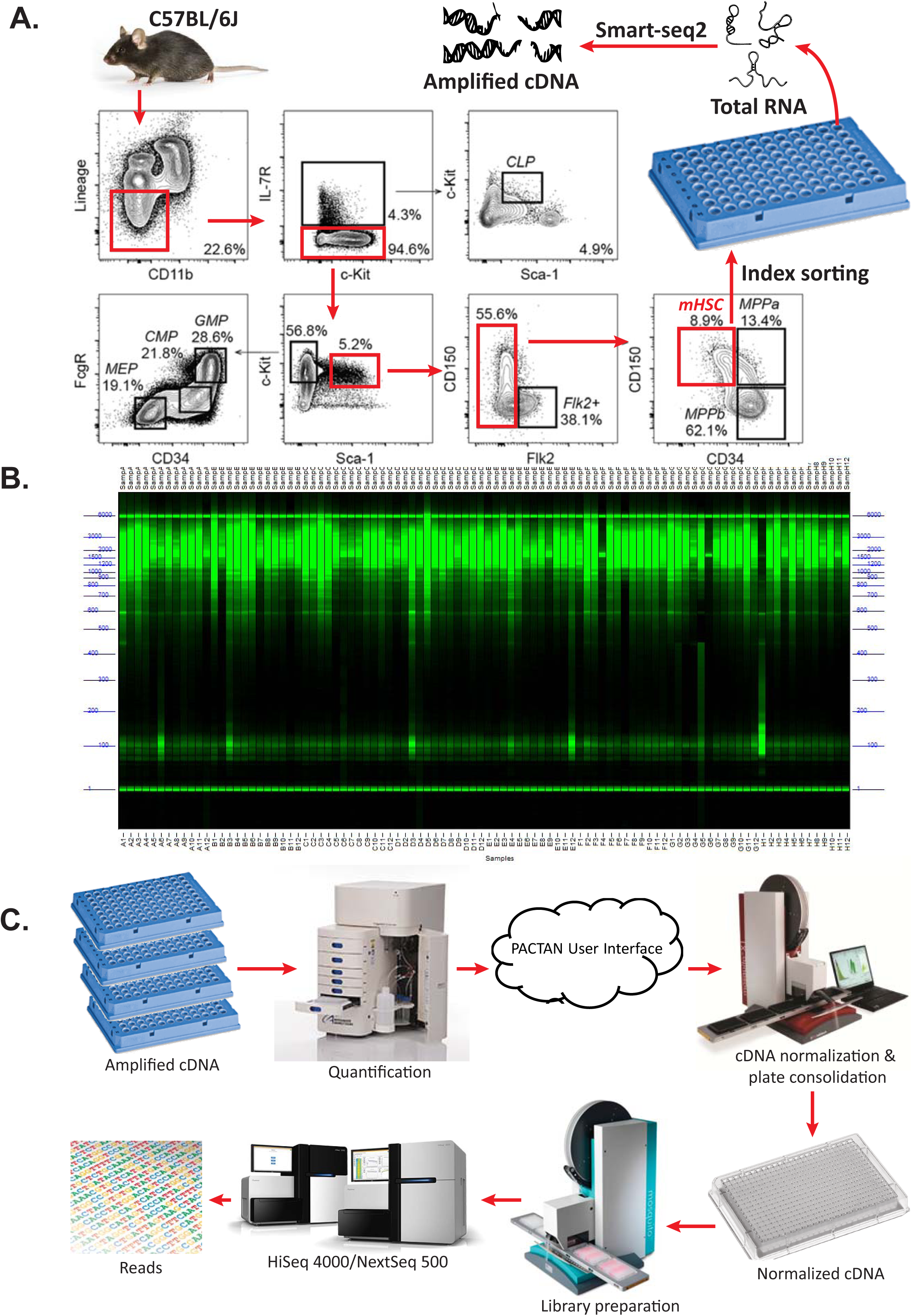
Modified Smart-seq2 pipeline. **A)** Schematic showing gating strategy^17^ on a FACS Aria II cell-sorter (BD Biosciences) to isolate mouse hematopoietic stem cells (mHSCs) and multipotent progenitors (mMPPs) as index-sorted single cells directly into 96-well PCR plates containing 4 μl of lysis buffer in each well. Poly-A mRNA in the cell lysate is converted to cDNA, which is then amplified using the Smart-seq2 protocol. **B)** Representative image of ethidium bromide-stained amplified cDNA smears from individual cells analyzed on a high-throughput Fragment Analyzer (AATI). **C)** Semi-automated downstream workflow. First, cDNA from single cells in individual wells of multiple 96-well plates are assessed for quality and quantity using the Fragment Analyzer in a high-throughput manner, providing values for each cDNA smear (size range 500 to 5000 base pairs), like that shown in B. The quantification data is then imported into our custom software, PACTAN (<u>p</u>rogram for <u>a</u>cquisition, <u>c</u>onsolidation, <u>t</u>racking, <u>a</u>nd <u>n</u>ormalization)^18^, which selects “good” wells with usable amounts of cDNA, calculates a dilution scheme to normalize cDNA across all individual wells and across all 96-well plates to obtain the desired range of cDNA concentration, develops a strategy to consolidate normalized cDNA into multiple 384-well plates as needed, and generates worklists compatible with liquid handling robots for downstream processing—all while tracking each well from sorting to sequencing. Using worklists generated by our software, liquid handling robots (Mantis from Formulatrix, Mosquito X1, and Mosquito HTS from TTP Labtech) perform all the pipetting steps to normalize and consolidate cDNA into 384-well plates, and prepare Nextera XT libraries. 384 individual libraries from each 384-well plate are then pooled. Finally, 384-plex pooled libraries are cleaned-up using AMPure XP beads and sequenced at our core facilities on a single lane of HiSeq 4000 (∼300 million clusters/384 cells) or NextSeq 500 (∼400 million clusters/384 cells).

## Results

### Spreading of signal on HiSeq 4000

In a single-cell RNA-seq experiment involving sorted mHSCs, we observed that signal from a highly-expressed gene in a given well (a single cell) was spread across wells that shared the same row or the column in the 384-well plate that was used to prepare libraries. We call this phenomenon “spreading-of-signal” **(Fig. 3)**.

**Figure 3.**
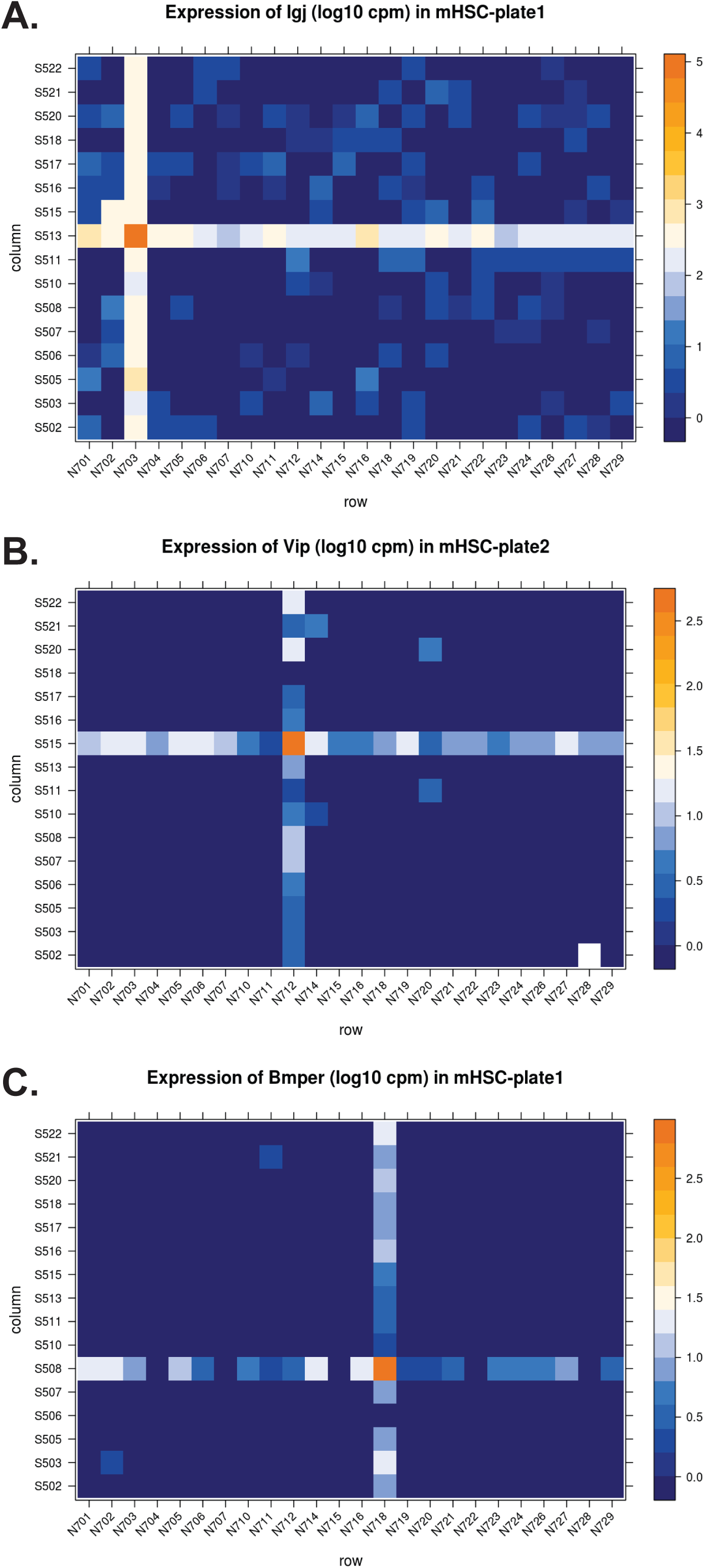
“Spreading-of-signal”. **A-C)** 384-well layout showing expression in log read counts per million (CPM) of the three representative genes indicated in a single-cell RNA-seq experiment where libraries from single mHSCs were prepared as in Figure 1 followed by sequencing on a HiSeq 4000. Reads were mapped to the mouse genome (mm10) and CPM values for each gene in each cell was calculated. CPM values for a given gene were then plotted in the 384-well layout such that each square shown on the 384-well layout corresponds to the expression value for the gene in the original cell from which the library was prepared.

Before we discovered this signal spreading, we had processed mHSC single cell RNA-seq data generated on the HiSeq 4000 with our previously published computational method^13^ that performs iterative robust principal component analysis (rPCA) to identify subpopulations of cells. In so doing, we identified 41 sub-populations of mHSCs, in line with our initial goal of identifying cell-intrinsic heterogeneity within a seemingly homogeneous mHSC population. However, upon further inspection of the data, we found that each of the single cells within each putative mHSC subpopulation mapped-back either to a single row or to a single column in the 384-well plate that was used to prepare the single cell libraries. We next inspected the expression levels of individual genes that drove classification of the 41 putative mHSC subpopulations. We found that if a given gene was expressed at a very high level in a well (that contained cDNA from a single cell) in the 384-well plate, the signal was also detected but at lower levels at all or most wells in the same row and column, forming a characteristic cross-like expression pattern in the plot **(Fig. 3)**. Since individual cells were initially placed at random in wells during single-cell-sorting and each library from a given cell was prepared independently, we inferred that this characteristic, cross-like pattern must have arisen from an artifact or systematic error that arose during or subsequent to sorting. This also implied that the 41 putative subpopulations of mHSCs identified by our subsequent computation analysis of the single cell expression data was not biologically meaningful, but was the result of the artifact or systematic technical error.

To identify the origin of the error, we performed the following three experiments.

### Experiment 1: signals in empty wells

We first took advantage of an ongoing single-cell RNA-seq experiment studying mouse fetal heart cells (mFHCs) where in a particular pool (“TYP4”) cDNA had been intentionally omitted from more than half the wells in the 384-well plate **(Fig. 4A)**. There were two kinds of empty wells: 1) wells in which no cDNA or reagents were added, and 2) wells in which no cDNA but all reagents, including index primers, were added. These latter wells had received the reagents from a 16-channel liquid handling robot, Mosquito HTS (TTP Labtech), which pipettes into 16 wells at once, column by column, while preparing libraries as per the Nextera XT protocol (Illumina). The bioanalyzer trace for this pool after cleanup is shown in **Fig. 4B.**

**Figure 4.**
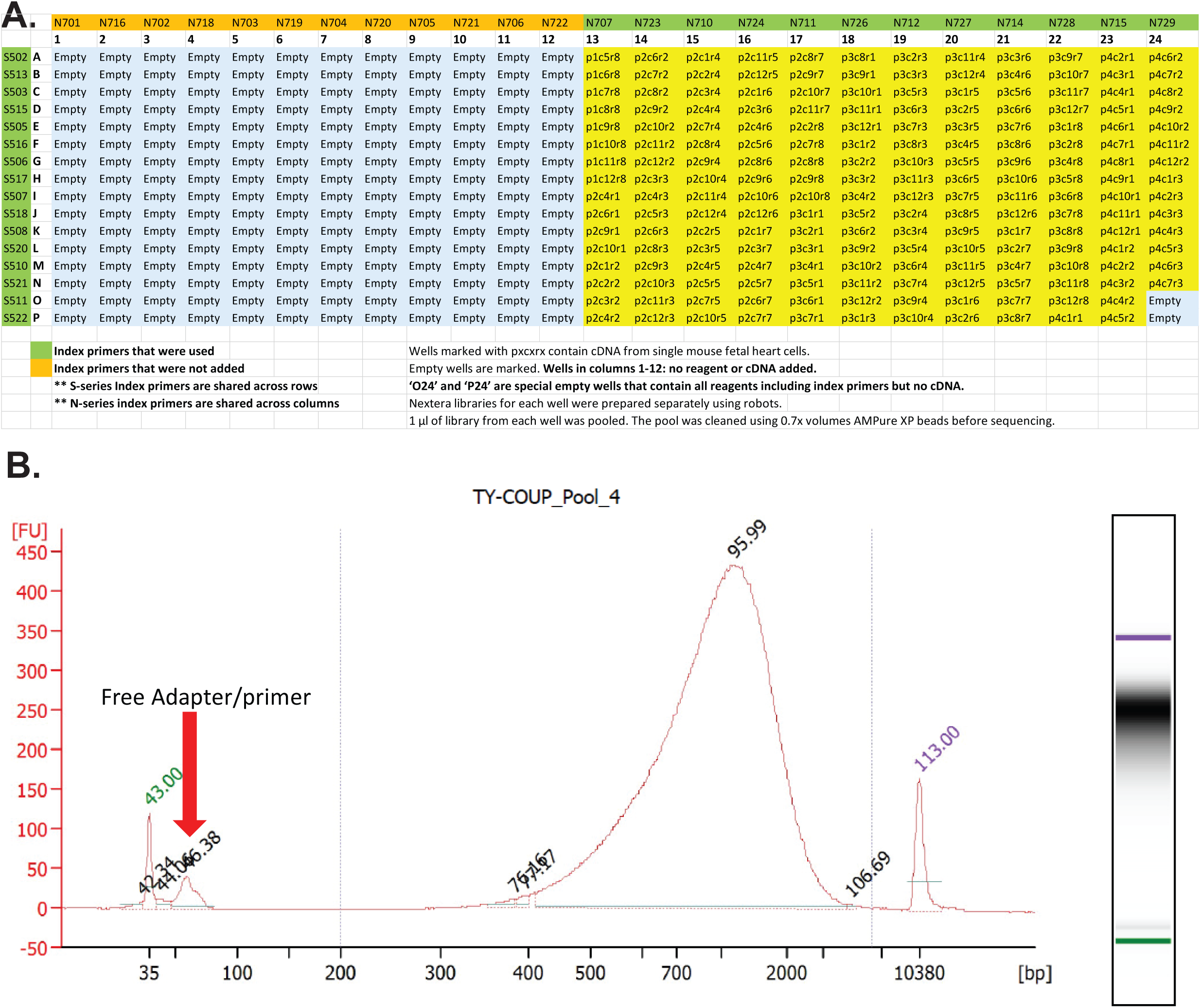
**A)** “TYP4” pool 384-well layout showing map of wells containing normalized cDNA from individual cells. **B)** After preparing libraries independently in each well in the columns (13-24) as shown in A, the individual libraries were pooled and cleaned-up using 0.7x AMPure XP beads. The cleaned library pool was analyzed on a Bioanalyzer (Agilent). Plot shows fragment distribution and concentration (peak-intensity) of the constituent library molecules and leftover free index primers after clean up (red arrow).

We found that, surprisingly, empty wells containing reagents (including index primers) but no cDNA had a significant number of high quality reads assigned to them. These reads mapped to the mouse genome (mm10) with ∼80% efficiency, similar to the efficiency of wells that contained cDNA **(Fig. 4A, 5, and 6)**. The number of mapped reads assigned to these empty wells was 5-7% of the average number of mapped reads assigned to wells that contained cDNA. In contrast, empty wells without cDNA or reagents had few reads assigned to them that passed our quality filter **(Fig. 4A, 5, and 6),** and only ∼50% of these reads mapped to the mouse genome. This phenomenon was consistent across many (n>50) 384-plex single-cell pools that were prepared by different researchers from different laboratories (n>8) at Stanford University.

**Figure 5.**
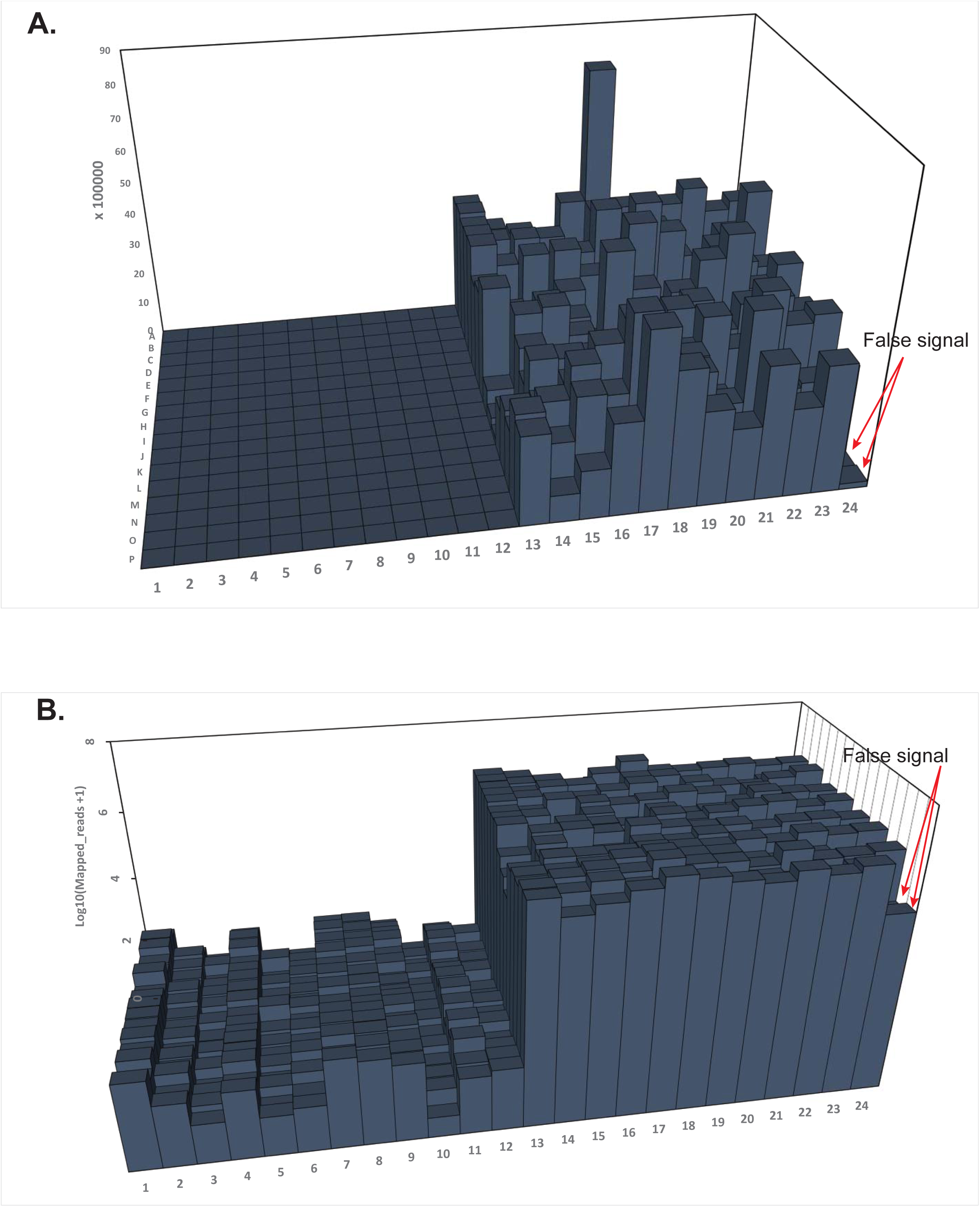
Bar-graph showing the number of mapped reads in **A)** linear and **B)** log scale resulting from individual libraries in the “TYP4” pool after sequencing on HiSeq 4000 at SFGF. Bars correspond to the 384-well layout as described in Figure 3A. Red arrows indicate false signals (above background signal) from wells O24 and P24 (as shown in Figure 3A).

**Figure 6.**
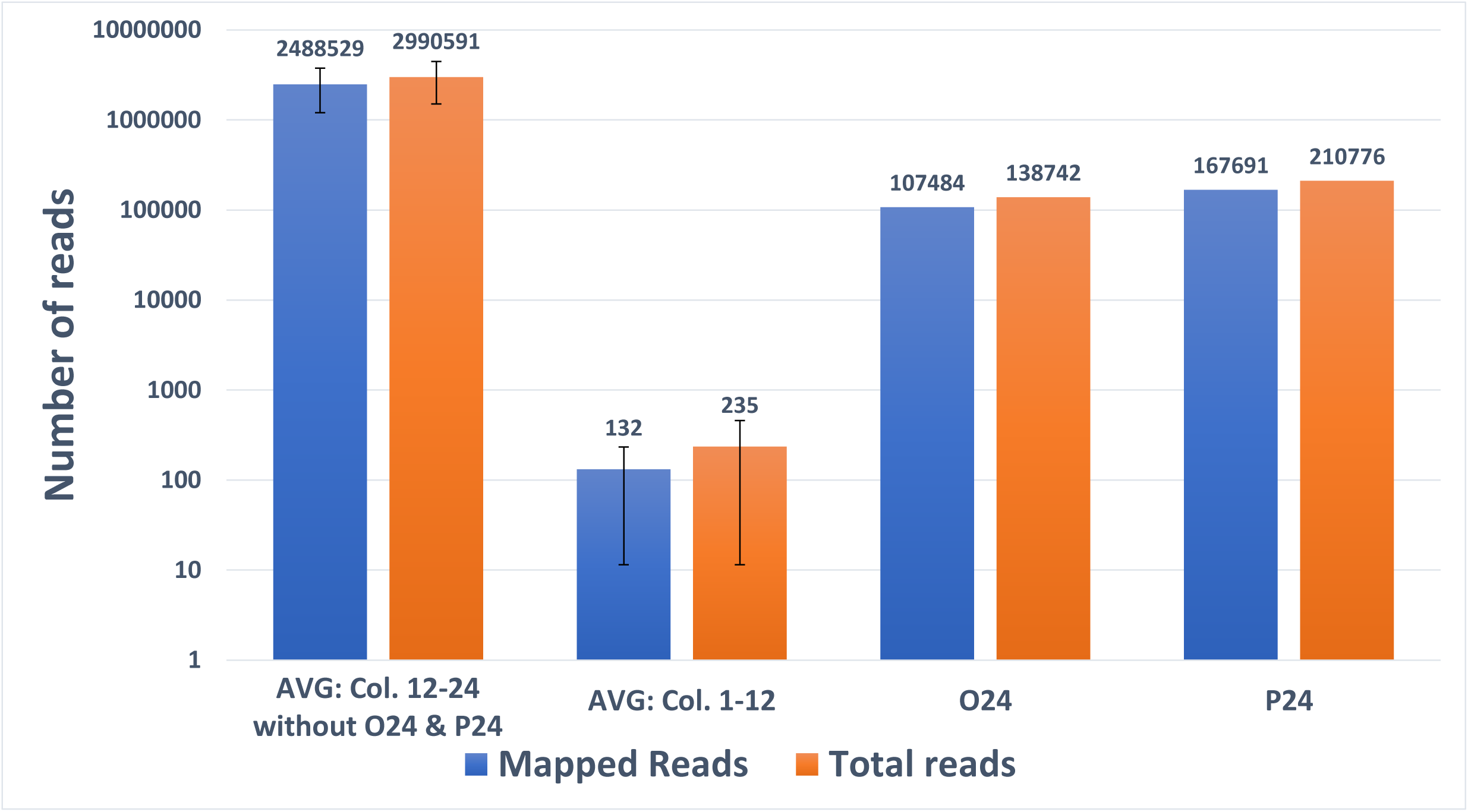
Plot showing the number of mapped reads (blue) and total reads (orange) for the different kinds of wells described for the “TYP4” pool in Figure 3A. Error bars, standard deviation of mean values.

These results suggested that the 5-7% of high-quality reads in the wells, ‘O24’ and ‘P24’ (containing only reagents and primers), originated from other wells in the pool that had cDNA. It further suggested that 5-7% of reads in all wells that had cDNA contain reads from other wells with cDNA, and this cross-contamination is caused by the presence of free index primers, because index primers were added to wells ‘O24’ and ‘P24’ but not to wells belonging to columns 1-12 **(Fig. 4A, 5, and 6).**

### Experiment 2: signals due to excess free index primers in a library pool

To further investigate the apparent cross-contamination, we added an excess amount of the index primers N701 and N702 into a leftover aliquot of single-cell pool “TYP4” (spike-in#1). The N701 and N702 index primers were used because they had not been added at any stage during library preparation in “Experiment 1” above **(Fig. 7A).** We then divided this “TYP4” spike-in#1 pool into two tubes and sequenced the contents of each tube separately on a HiSeq 4000 and a NextSeq 500 at the Stanford Functional Genomics Facility (SFGF). Surprisingly, after demultiplexing, data from the HiSeq 4000 had a very large number (up to ∼7 million) of high-quality reads that were now assigned to all the wells in columns 1 and 3—the wells in the 384-well plate where no cDNA or reagents were added. Furthermore, a large number of these reads (up to 3.5 million) now assigned to wells in columns 1 and 3 mapped to the mouse genome (∼50% mapping efficiency). Simply by adding excess free index primers (N701 and N702) not originally present in the “TYP4” pool, we obtained a substantial number of reads assigned to wells in column 1 and 3, as if we had added cDNA and reagents in these wells in the original “Experiment 1” above **(Fig. 8A, and B)**. In contrast, the same spiked-in#1 pool when sequenced on NextSeq 500 resulted in significantly fewer reads—on average 74 out of 116 total reads—mapping to the wells in columns 1 and 3, despite the presence of high levels of N701 and N702 index primers in the library pool **(Fig. 9A, and B).**

**Figure 7.**
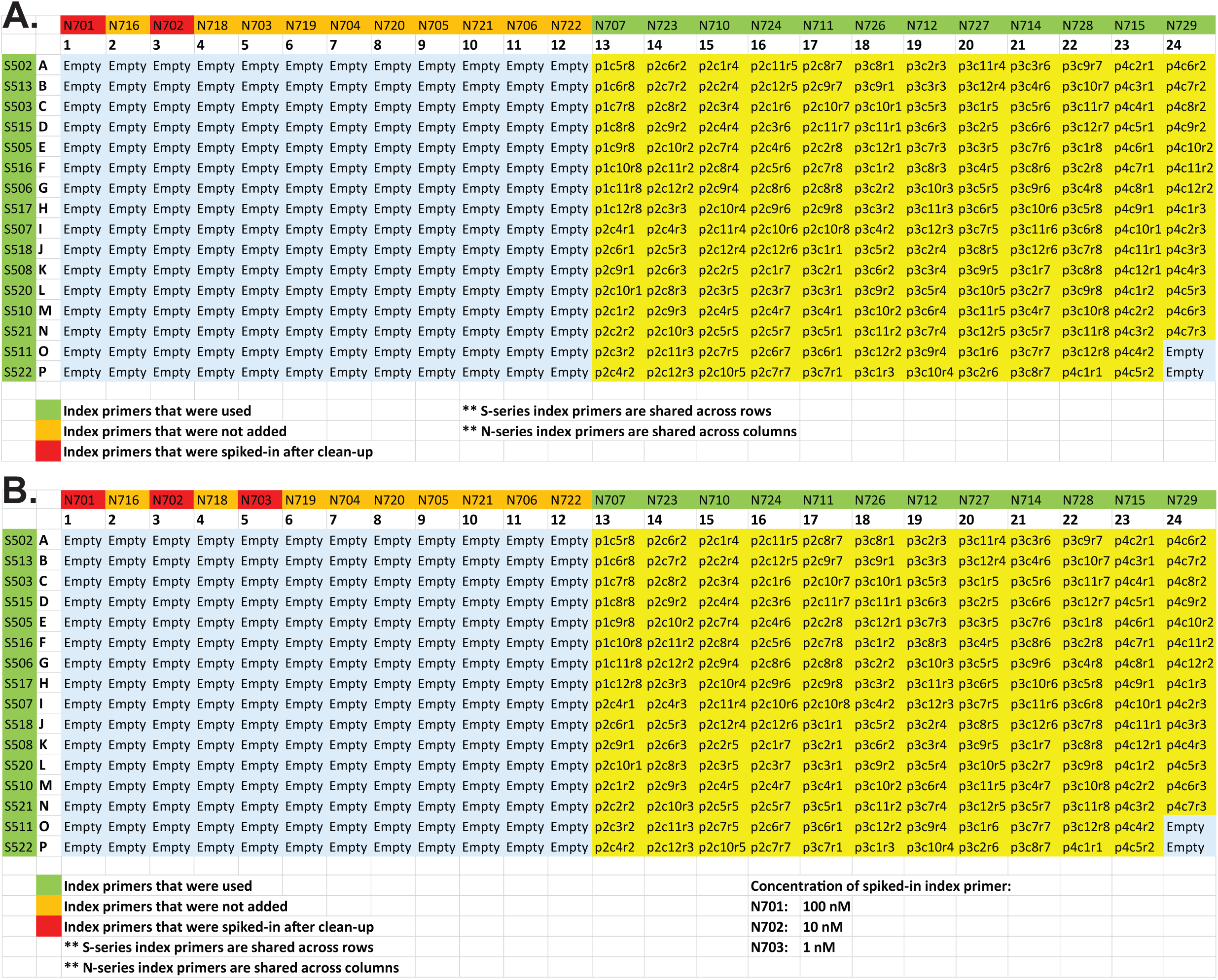
Diagram showing layout of “TYP4-spike-in#1” (Experiment 2, **A)** and “TYP4-spike-in#2” (Experiment 3, **B)** superimposed on a 384-well plate.

**Figure 8.**
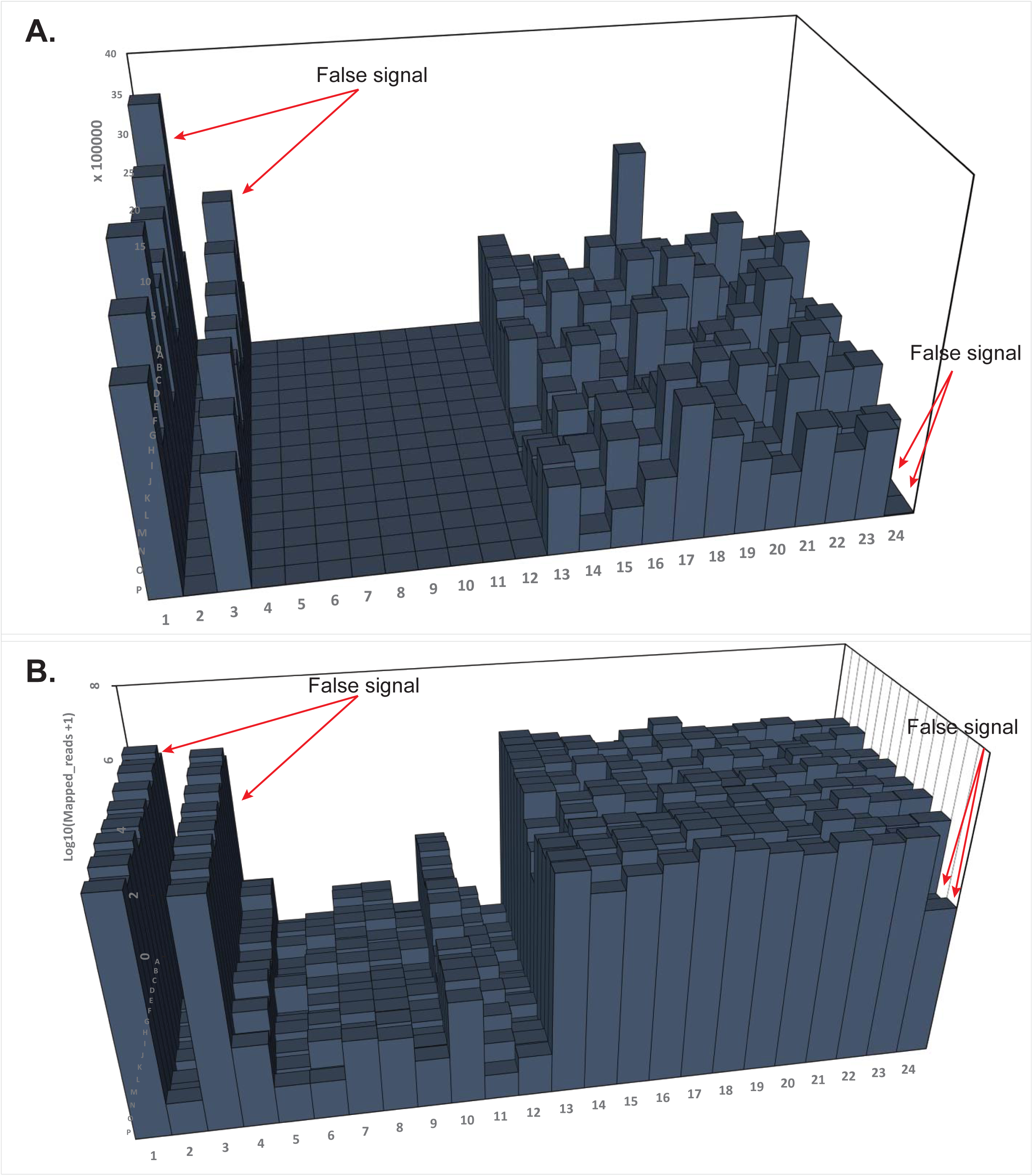
Plot showing number of mapped reads in **A)** linear and **B)** log scale resulting from individual libraries in the “TYP4-spike-in#1” pool (Experiment 2) after sequencing on HiSeq 4000 at SFGF. Values (bars) are shown in the 384-well layout as described in Figure 6A. Red arrows, false signals (above background signal).

**Figure 9.**
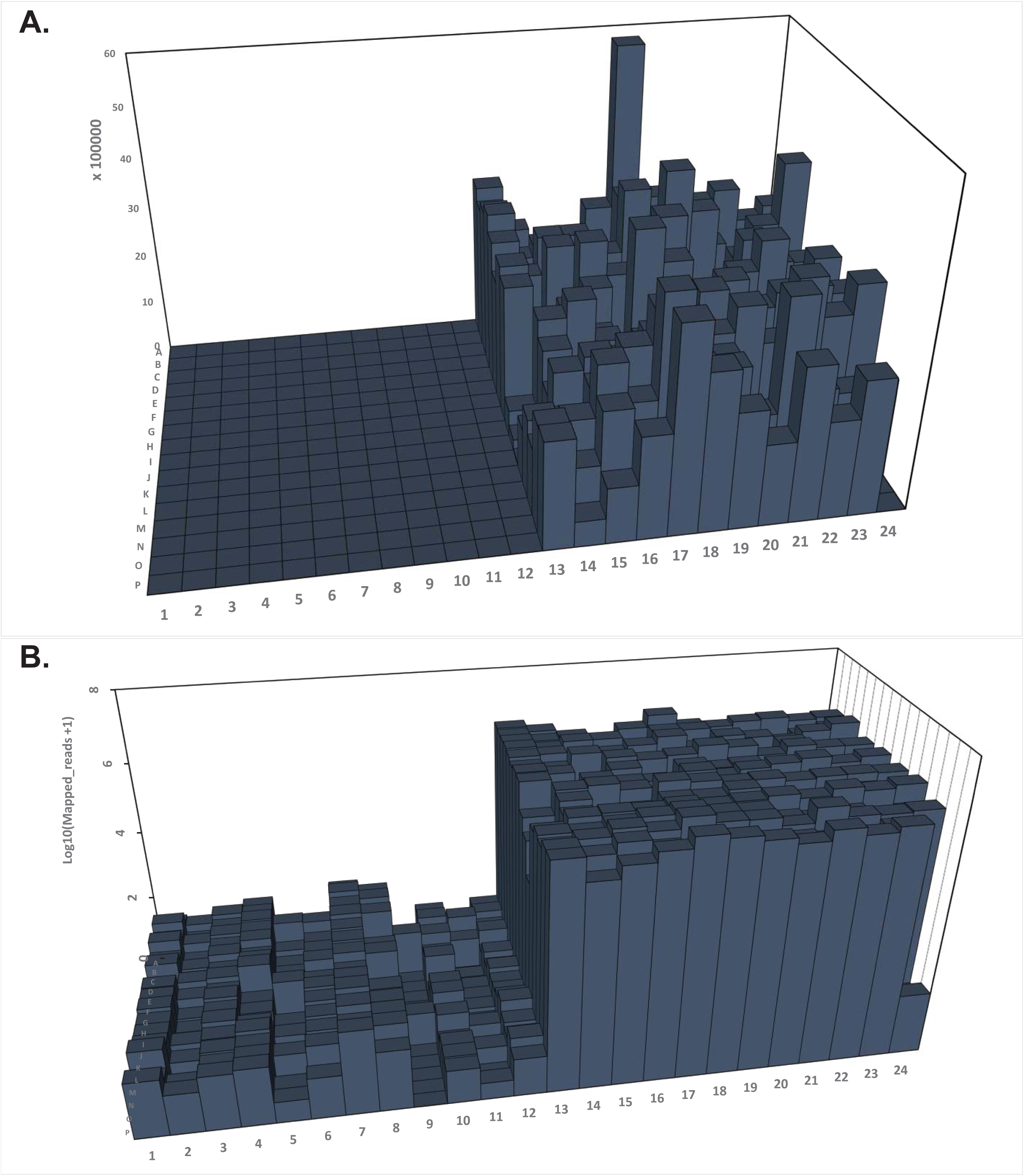
Plot as in Figure 7 showing number of mapped reads in **A)** linear and **B)** log scale resulting from individual libraries in the “TYP4-spike-in#1” pool (Experiment 2) after sequencing on NextSeq 500 at SFGF.

### Experiment 3: spreading-of-signals on a different HiSeq 4000 instrument

We wondered if the observed effect was specific to the machines or methods for sequencing at the SFGF core facility, or were more general. To test this, we sought to independently verify the SFGF results. We therefore took an additional aliquot of the “TYP4” pool from “Experiment 1” above and added Illumina index primers N701, N702, and N703 at concentrations 100 nM, 10 nM, and 1 nM, respectively (spike-in #2 pool) **(Fig. 7B).** As in Experiment 2 above, we divided the TYP4-spike-in#2 pool in half and sequenced the first half on a HiSeq 4000 at the Stanford Center for Genomics and Personalized Medicine (SCGPM), and the other half on the NextSeq 500 in the Stephen Quake lab. As in “Experiment 2”, when spike-in#2 pool was sequenced on a HiSeq 4000 at SCGPM we found that index switching resulted in high-quality reads that were assigned to all wells in columns 1, 3, and 5, which had no cDNA or reagent added in experiment 1 **(Fig. 10A, and B).** Further, the number of mapped reads assigned to these columns was directly proportional to the amount of spiked-in index primer. By contrast, sequencing of the same spike-in#2 pool on the Quake lab NextSeq 500 did not show large-scale index switching **(Fig. 11A, and B).**

**Figure 10.**
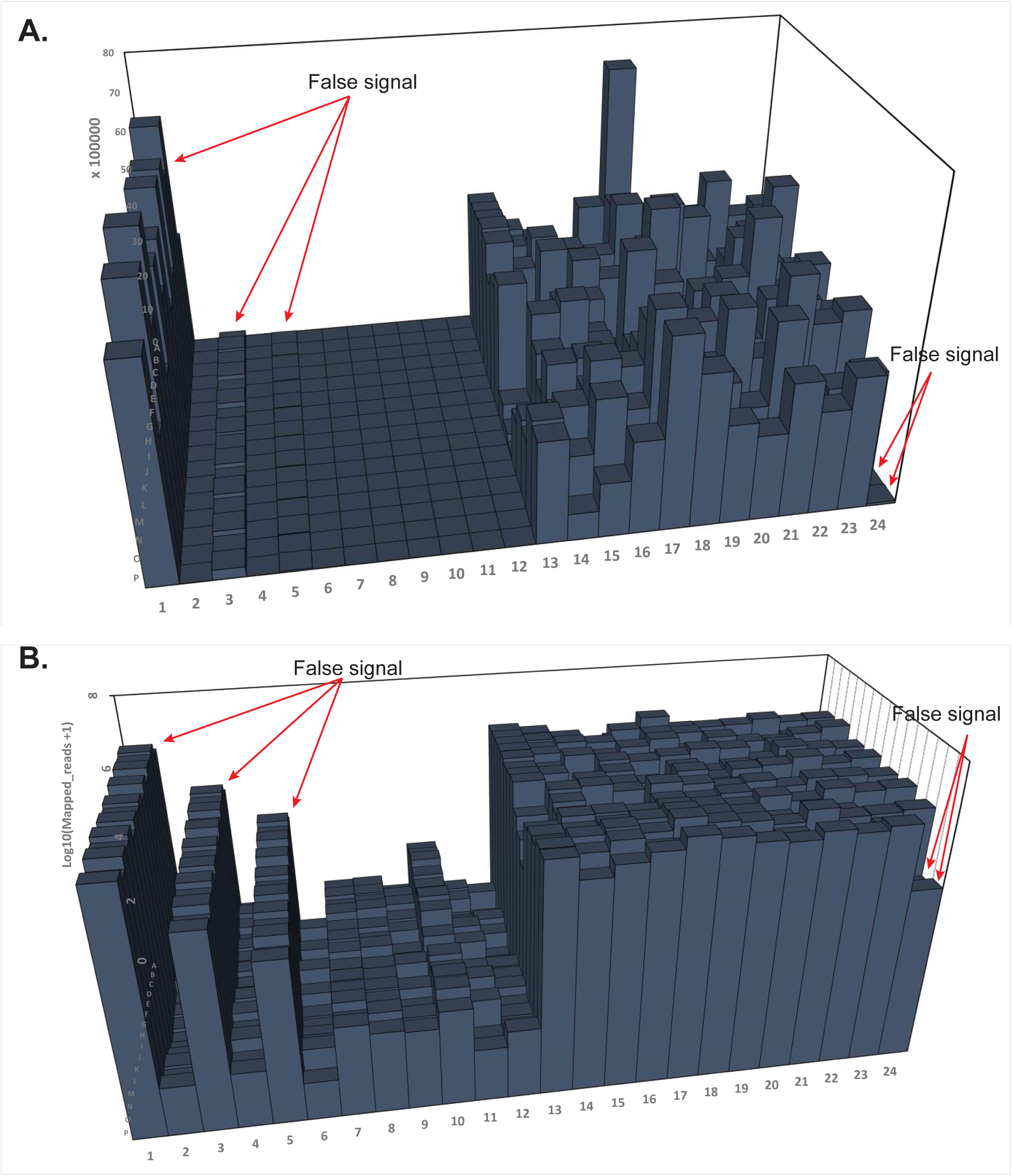
Plot showing number of mapped reads in **A)** linear and **B)** log scale resulting from individual libraries in the “TYP4-spike-in#2” pool (Experiment 3) after sequencing on HiSeq 4000 at SCGPM. Values (bars) are shown in the 384-well layout as described in ‘Figure 6B’. Red arrows, false signals (above the background signal).

**Figure 11.**
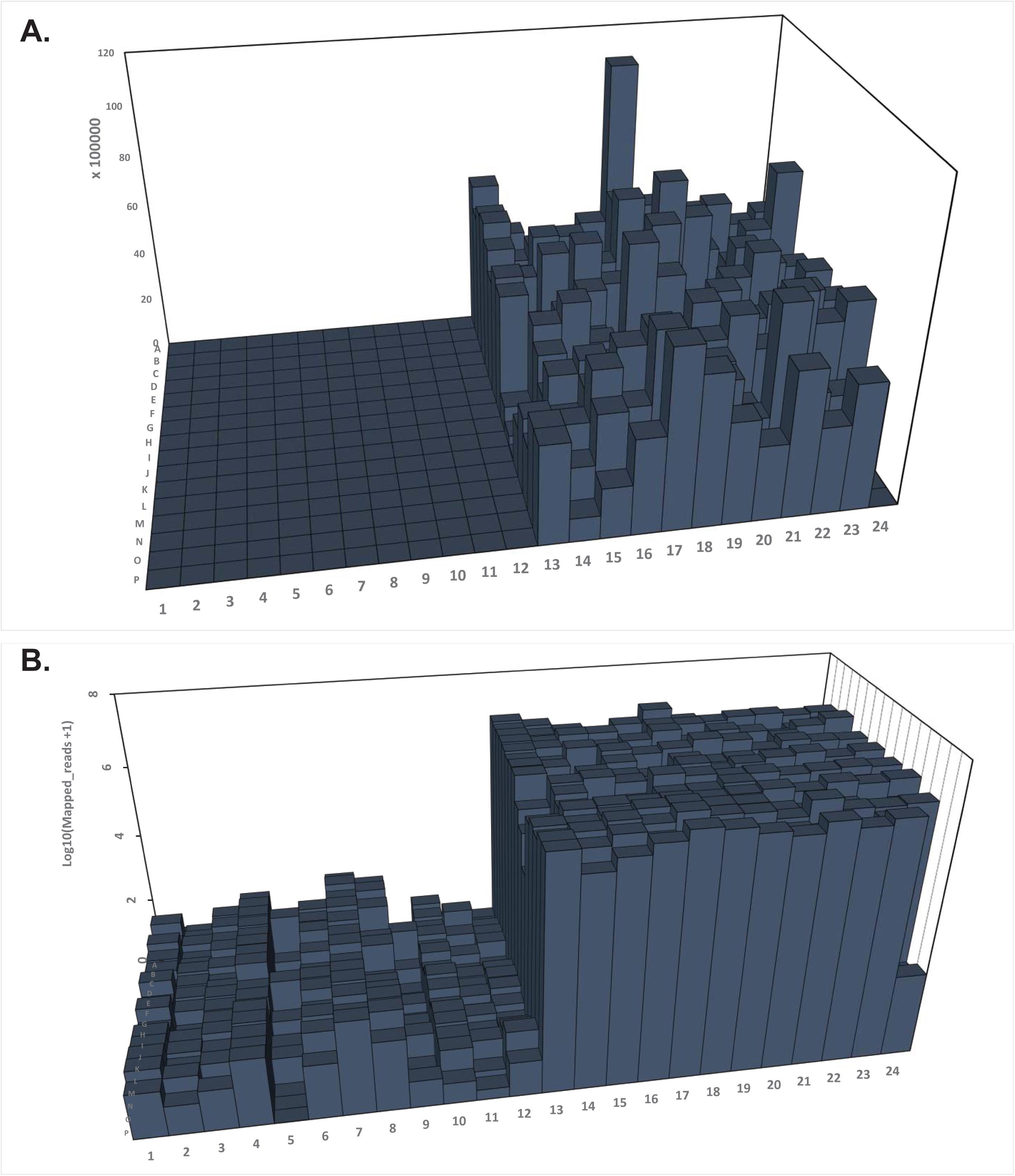
Plot as in Figure 9 showing number of mapped reads in **A)** linear and **B)** log scale resulting from individual libraries in the “TYP4-spike-in#2” pool (Experiment 3) after sequencing on NextSeq 500 at Stephen Quake laboratory.

### Spreading of signals HiSeq 4000 vs NextSeq 500

After demonstrating that index switching occurs in the presence of free index primers, we revisited our mHSC experiment where we had first observed the “spreading-of-signal” across rows and columns. Resequencing of these pools on a NextSeq 500 showed that the “spreading-of-signal” effect observed when DNA sequences were obtained with a HiSeq 4000 **(Fig. 3)** was no longer present when sequences were obtained with a NextSeq 500 **(Fig. 12)**. Multiple labs at Stanford (n>4) have similarly resolved the “spreading-of-signal” issue observed using a HiSeq 4000 by resequencing their single-cell RNA-seq libraries on a NextSeq 500.

**Figure 12.**
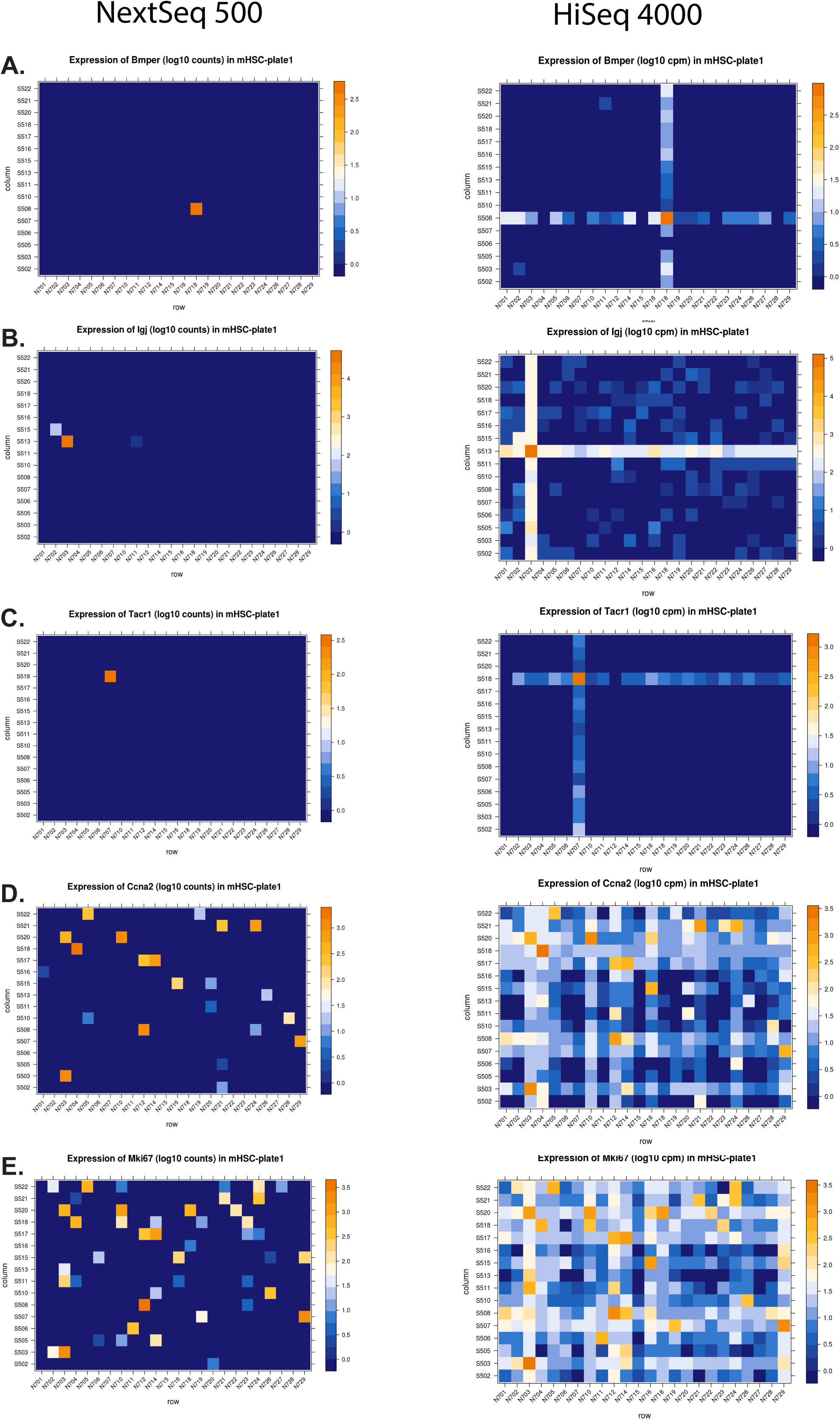
Spreading-of-signal: comparison between HiSeq 4000 and NextSeq 500. **A-E)** 384-well layout showing expression in log CPM of the five representative genes indicated in the same mHSC single-cell RNA-seq experiment as shown in Figure 2. Reads resulting from HiSeq 4000 or NextSeq 500 were mapped to the mouse genome (mm10) and CPM values for each gene in each cell were calculated. The CPM values for a given gene are plotted in a 384-well layout such that each square in the layout corresponds to the expression value of the gene in the original cell from which the library was prepared.

## Discussion

We conclude that the free index primer levels remaining in our library pools after a stringent cleanup procedure (0.6-0.7x volume AMPure XP beads) resulted in 5-7% of reads in each sample possibly belonging to other samples in the library pool, most likely due to index switching at random. Since the HiSeq 3000 and HiSeq X Ten share the same chemistry as the HiSeq 4000, it is possible that such index switching may also occur at a similar rate using these sequencers, although we have not tested this directly.

The systematic pattern with which this cross-contamination artifact affected our data was surprising and concerning. In our initial mHSC single-cell RNA-seq experiments described above, we found two putative major subpopulations, which we called mHSCa and mHSCb. Iterative rPCA analysis^13^ of each mHSCa and mHSCb identified 16 putative subpopulations of mHSCa, and 25 subpopulations of mHSCb. As noted above, cells in each apparent subpopulation arose from either a single column or a single row on the 384-well plate in which the individual libraries were prepared. There were 16 putative mHSCa subpopulations—which correspond to 16 S5xx index primers, and 25 mHSCb subpopulations—which correspond to 24 N7xx index primers. The 25^th^ mHSCb subpopulation was simply the leftover cells after the iterative rPCA analysis. Remarkably, all of the putative subpopulations identified were the result of the “spreading of signal” artifact, which results in the homogenization of expression signal among cells positioned in a row or column in the 384-well plate originally used to prepare the library.

The “spreading of signal” only among cells positioned in a given row or column in the 384-well plate most likely results from the conditions for isothermal amplification of library pool molecules on the cBot 2 system (Illumina)—the machine used to generate clusters on a patterned flow cell. The limited time allowed for the amplification reaction to take place in the cBot 2 during cluster generation (before sequencing) presumably restricts index switching to only one end of the most abundant library fragments (cDNA signals). This occurs when a free index primer—either forward (P7) or reverse (P5)— primes the library fragment via its complementary 3’ end and gets extended and thereby tags the newly synthesized library fragment with a new forward (P7) or reverse (P5) index **(Fig. 1)**. The probability of both indices switching in this manner is low, given the short time interval between cluster generation on the cBot 2 and initiation of sequencing on the HiSeq 4000. This would require a second priming event during this interval by another free index primer at the other end of the newly synthesized library fragment followed by extension, prior to library fragment binding to the patterned flow cell.

In our dual index setup described above, an index swap at one end of a library results in signal spread among samples (wells) along the same row or column in the plate, which is what we commonly observed. Index swapping at both ends would result in spread of signal among samples (wells) throughout all rows and columns in the plate; this occurs too, albeit at a much lower frequency **(Fig. 12B)**.

The “spreading-of-signal” was not limited to single-cell RNA-seq libraries, but was common to all library pools (bulk RNA-seq, ATAC-seq etc.) in which samples were multiplexed and some level of free index primers were present. Once we understood how to detect the index switching, it was easy to discern whether a given dataset was affected by this issue: when de-multiplexing, we simply looked for reads carrying a set of unique Nextera forward and reverse index combinations that normally would not be expected to have reads assigned to them, similar to empty wells with reagents as shown above in Experiments 1, 2, and 3 above.

However, it remains difficult to detect the level of cross-contamination in single index libraries, libraries in which only one index is present on one side of the DNA fragment to be sequenced, unless PhiX control (Illumina) was added to the library pool before sequencing. If PhiX was added, then index switching resulted in a significant number of PhiX molecules acquiring all free indices that were used in a given multiplexed library pool (data not shown).

Nonetheless, caution is warranted in interpreting such data, especially when the exact adapter sequences at the ends of PhiX control library molecules are not publically disclosed by Illumina. The PhiX control supplied by Illumina is non-indexed to begin with, and therefore, clusters that result from such PhiX molecules are “dark”, i.e. there is no fluorescent signal coming from the PhiX clusters when the indices are being read by the sequencer. Therefore, index assignment to PhiX reads could arise for three different reasons: 1) background sequencing error, 2) indices falsely assigned to a PhiX cluster if it was in close proximity to another cluster with index on the flow cell, and 3) the ExAmp related index-switching described above.

There are third party suppliers that offer indexed PhiX libraries, which should minimize index mis-assignment resulting from the first two reasons described above. Our center is switching to an indexed PhiX spike-in scheme, which will let us more accurately determine the amount of index switching that occurs exclusively as a result of the ExAmp chemistry in a given multiplexed library pool on the HiSeq 4000.

## Conclusions

Overall, across multiple single-cell experiments, we estimate that the level of cross-contamination of assignments in multiplexed samples is between 5-10% of total mapped reads, the value depending on the level of free index primers present in a library pool, and therefore, variable among experiments. We also want to emphasize that, in an experiment, if different samples with different indices are not multiplexed and only a single sample carrying only one index (or a unique set of two indices) is sequenced in a lane of a HiSeq 4000, we would not expect “spreading-of-signal”.

Our results suggest multiple workaround strategies that could mitigate or even eliminate this issue in the newer sequencers with patterned flow cells. For example, 1) creating bar code sets that do not use overlapping pairs, or, 2) developing more thorough cleanup approaches for removing free primers from library pools, for example, uracil-DNA glycosylase (UDG) digestion^14-16^, or polyacrylamide gel electrophoresis (PAGE) based purification of library pools. PAGE-purification of library pools has diminished over the past 4-5 years in favor of purifications using AMPure XP beads, primarily because of the ease-of-use of beads and the potential for automation using liquid handling robots.

The experiments described here likely represent only a tiny fraction of experiments at Stanford and worldwide whose analysis and conclusions are compromised by the described “spreading of signal” artifact. The toll includes highly precious and hard-to-isolate normal and irreplaceable patient samples, time devoted to the performance and analysis of such compromised experiments by pre- and postdoctoral researchers on a limited clock for training, and funds from federal and state government and philanthropic agencies for advancing medical science and developing new diagnostics and therapies.

## Acknowledgements

We thank Stephen Quake, Norma Neff, Seth Karten and John Coller for valuable suggestions and critical reading of the manuscript. We further thank SFGF (Director: John Coller) and Norma Neff for the help in the design and execution of “Experiment 2” and “Experiment 3”, respectively. The data generated on the Illumina HiSeq 4000 machines in SFGF and SCGPM was purchased with funds from NIH under award numbers S10OD018220 and 1S10OD02014101. Funding for the research came from NIH grant R01CA086065, and CESCG/CIRM grant GC1R-06673-A to ILW, and Ludwig Cancer Foundation.

---

**Supplementary Figure 1.** The ExAmp procedure for cluster generation in the HiSeq 4000^3,6^. Single stranded library molecules—resulting from the alkali denaturation of double-stranded library molecules—are mixed with the ExAmp clustering reagents (Illumina) and loaded onto a patterned flow cell. The flow cell is tiled with billions of nanowells, where each nanowell surface is tiles with a lawn of P5 and P7 primers^7^. The movement and seeding of library molecules into nanowells is controlled by an electric field, that guides library molecules to nanowells while they are repelled from the interstitial space between nanowells. The electric field is also used to control the diffusion rate of library molecule. For effective cluster generation, the rate of diffusion that leads to seeding of library molecules in the nanowells must be slower than the rate of isothermal amplification that generates a cluster. This helps ensure that a single-stranded library molecule gets amplified rapidly and generates a cluster before another library molecule seeds the same nanowell. However, if more than one library molecule does seed the same nanowell, a “polyclonal” cluster can result, thus leading to a drop in base call quality while sequencing. If the loading concentration of a library pool is optimal, seeding of nanowells should follow a Poisson distribution with most nanowells seeded by one or no library molecules, and only a few nanowells seeded by multiple library molecules to generate polyclonal clusters.

If multiple library molecules seed a nanowell, a different rate of amplification for each library molecule increases the chance that only one library molecule will result in a cluster. This is achieved through a strategy in which the rate at which the first amplicon is synthesized during the first round of amplification is relatively slower than the subsequent amplification steps following the first one. The ExAmp reagent cocktail (Illumina) contains capture primers. The 3’ ends of these primers are blocked by a dideoxy nucleotide. These primers anneal to ends of library molecules thereby blocking the seeding of library molecules in a nanowell. In the presence of excess pyrophosphate, the polymerase removes the dideoxy block from the 3’ end of the capture primer^19-22^. Certain (proprietary) conditions ensure that on average only one capture primer in a nanowell gets unblocked and extended by polymerase, even if multiple capture primers are bound to different library molecules. The newly synthesized complementary strand of the library molecule is now separated from the original template aided by single-strand-binding proteins and helicases and anneals to a P5 or a P7 primers that are physically linked to the flow cell. Since the 3’ ends of P5 and P7 primers contain no dideoxy nucleotide block the annealed complemtary strand of the library molecule and can undergo rapid amplification via isothermal bridge amplification to form a cluster. The isothermal bridge amplification is much more rapid than the stochastic removal of dideoxy-block from the 3’ end of the capture primer and its extension. Therefore, before other capture primers in the nanowell bound to other library molecules are converted to an extensible state, the library molecule—where the first capture primer gets unblocked and extended—binds the P5 or P7 primer forming an amplification site and kinetically excludes and outcompetes other library molecules for cluster generation. Compared to the older method, the resulting clusters can be at a much higher density per unit area of the flow cell, because each cluster is confined within a clearly demarcated nanowell. This also mitigates the loss in quality that can result from overloading a flow cell. Moreover, a different metric (compared to the older method) is used to assess the quality of these clusters, filtering out dim (unseeded nanowells), low quality, and polyclonal clusters during sequencing, further enhancing read quality.

**Supplementary Figure 2.**
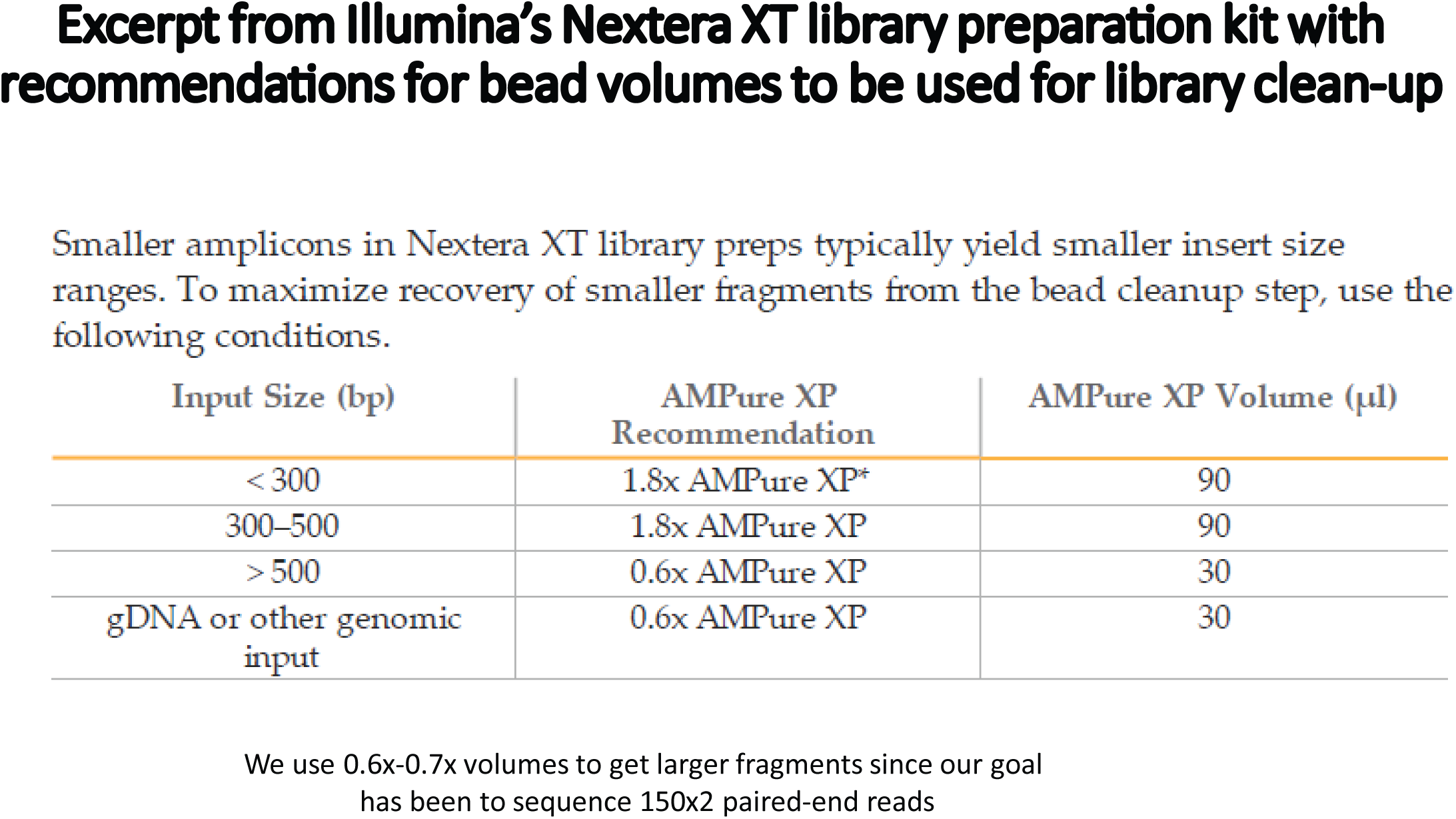
Image of the Nextera XT protocol (Illumina) showing AMPure XP bead volumes recommended for library pool cleanup.

